# UniWave: A Waveform-Based Encoding Framework for Nucleic Acid Feature Extraction

**DOI:** 10.64898/2026.01.12.698567

**Authors:** Hao Zhang, Yujun Qi, Lili Wang

## Abstract

Nucleic acid sequence analysis constitutes a core research area in biomedical and health informatics, playing a critical role in infectious disease surveillance, epigenetic regulation, and genomic biomarker discovery. However, most existing sequence encoding methods rely on discrete representations, which are inadequate for capturing the intrinsic continuous structural properties of biological sequences. Inspired by waveform representations in modern physics, we propose UniWave, a novel encoding framework that converts discrete nucleotide sequences into biologically informative one-dimensional continuous waveforms through base mapping, windowed sinc interpolation, and wavelet-based downsampling. To enable more efficient spatiotemporal modeling, we design a learnable positional encoding module, WavePosition, which incorporates positional information to project the one-dimensional waveform into a two-dimensional continuous representation. In addition, we develop a lightweight dual-attention network, InceptionTime-ATT, to facilitate efficient multi-scale extraction of biological features. Experiments conducted on multiple representative genomic benchmark tasks demonstrate that UniWave consistently and significantly outperforms conventional encoding methods in viral genotype classification, cross-species enhancer identification, epigenetic modification site detection, and bacterial genome classification. Moreover, UniWave maintains excellent robustness under perturbation and noise stress tests, while its low-dimensional continuous representation further contributes to reduced model complexity. In summary, UniWave establishes a novel paradigm for biomedical nucleic acid sequence encoding that is interpretable, noise-resilient, and computationally efficient, offering new avenues for uncovering disease-associated genomic patterns and advancing data-driven biomedical research.

## 1. Introduction

The increasing integration of computational approaches, including artificial intelligence and data mining, into biological research represents a defining trend for the future of the life sciences. Within this paradigm, nucleic acid sequence analysis has emerged as a cornerstone of biomedical and health informatics, underpinning a wide array of critical applications ranging from pathogen surveillance to epigenetic regulation. Driving these applications forward is feature extraction, which serves as a crucial bridge connecting raw biological data with computational models. Its primary function is to convert the genetic information embedded within biological sequences into machine-interpretable mathematical representations, thereby fundamentally setting the performance ceiling for downstream bioinformatics tasks. This process must preserve biologically meaningful information, including gene functional elements and genetic implications, while adhering to the mathematical structures required by computational models. Feature extraction goes beyond the mere conversion of “symbols” to “numbers”; it involves the systematic abstraction of multi-layered features encoded in gene sequences, encompassing local sequence motifs, long-range dependencies, and physicochemical properties. The core challenge lies in balancing biological interpretability, computational efficiency, and model generalizability.

Conventional feature engineering approaches predominantly depend on discrete modeling and domain-specific knowledge [1]. One-hot encoding [2, 3] provides a simple representation but suffers from high dimensionality and limited ability to capture interactions between nucleotides. K-mer-based features [4, 5] can represent local sequence patterns but are highly sensitive to the selection of k and are inadequate for modeling long-range dependencies. Physicochemical feature representations (e.g., PseNC) [6] can enrich sequence encoding, yet they rely on manually defined parameters and exhibit limited generalizability. Overall, these static approaches are unable to capture long-range interactions and are vulnerable to sequencing noise.

Deep learning facilitates end-to-end feature learning and the integration of complex biological signals. Convolutional neural networks (CNN) capture local features through multi-scale convolutions [7], whereas Transformers model global dependencies using self-attention mechanisms [8]. Self-supervised approaches, such as masked language modeling [9, 10], enhance generalization by leveraging unlabeled data. Nevertheless, interpretability remains a major challenge, and models like AlphaFold3 [11] require substantial computational resources, restricting their applicability in resource-limited environments.

To address the limitations of both conventional feature engineering and deep learning-based methods, we propose a novel feature encoding paradigm based on dynamic waveform modeling. From the standpoint of physics and mathematics, waveforms constitute the fundamental representation of dynamic processes. The propagation of biological genetic information can, in essence, be regarded as an evolving waveform process. Structurally, nucleotide sequences can be viewed as discrete fluctuations of A, T, C, and G nucleotides within a four-dimensional space. Consequently, genetic information inherently exhibits oscillatory characteristics, and converting sequence information into waveform signals enables a more intuitive representation of structural features. The linear organization of nucleotides encodes latent rhythmic and periodic fluctuation patterns: high-frequency short repeats correspond to waveform periodicities, mutation events manifest as localized perturbations, and long-range dependencies emerge as low-frequency components. These analogies highlight the potential of waveform-based representations to uncover structural and functional patterns within genetic sequences. In particular, multiscale waveform analysis techniques, including Fourier transforms and wavelet analysis [12, 13], provide a rigorous mathematical foundation for sequence feature extraction [14, 15] and hold significant promise for bioinformatics applications.

This study introduces the UniWave encoding framework as a novel contribution. The framework first establishes a mathematical relationship between nucleotide physicochemical properties and waveform representations through numerical mapping of physicochemical attributes. It subsequently generates a low-dimensional, high-information-density continuous waveform by integrating windowed Sinc interpolation, multiscale wavelet decomposition, and adaptive threshold-based downsampling. We innovatively integrate a learnable positional encoding module with a lightweight multi-scale hybrid attention network, enabling dynamic modeling of both local sequence features and long-range global dependencies. By interpreting biological information from a wave-based perspective, this approach facilitates the identification of latent regulatory patterns. Meanwhile, UniWave establishes a novel methodological foundation for genomic sequence analysis, and its outstanding performance across multiple core tasks provides validation of its technical feasibility and application potential. Building upon the present study, future work will focus on expanding the scope of applications, with particular emphasis on complex biological scenarios such as cross-species regulatory element prediction and cancer mutation detection.

## 2. Methods

This study presents UniWave, a dynamic waveform-based encoding framework for nucleic acid sequences, consisting of three core modules: data preprocessing, waveform encoding, and computational modeling. Collectively, these modules facilitate a systematic transformation of discrete sequences into continuous wave fields (Fig. 1). Following the complete encoding process, the generated data are stored in the standard HDF5 file format, enabling subsequent model training and evaluation.The subsequent sections provide a detailed description of each module.

**Figure 1.**
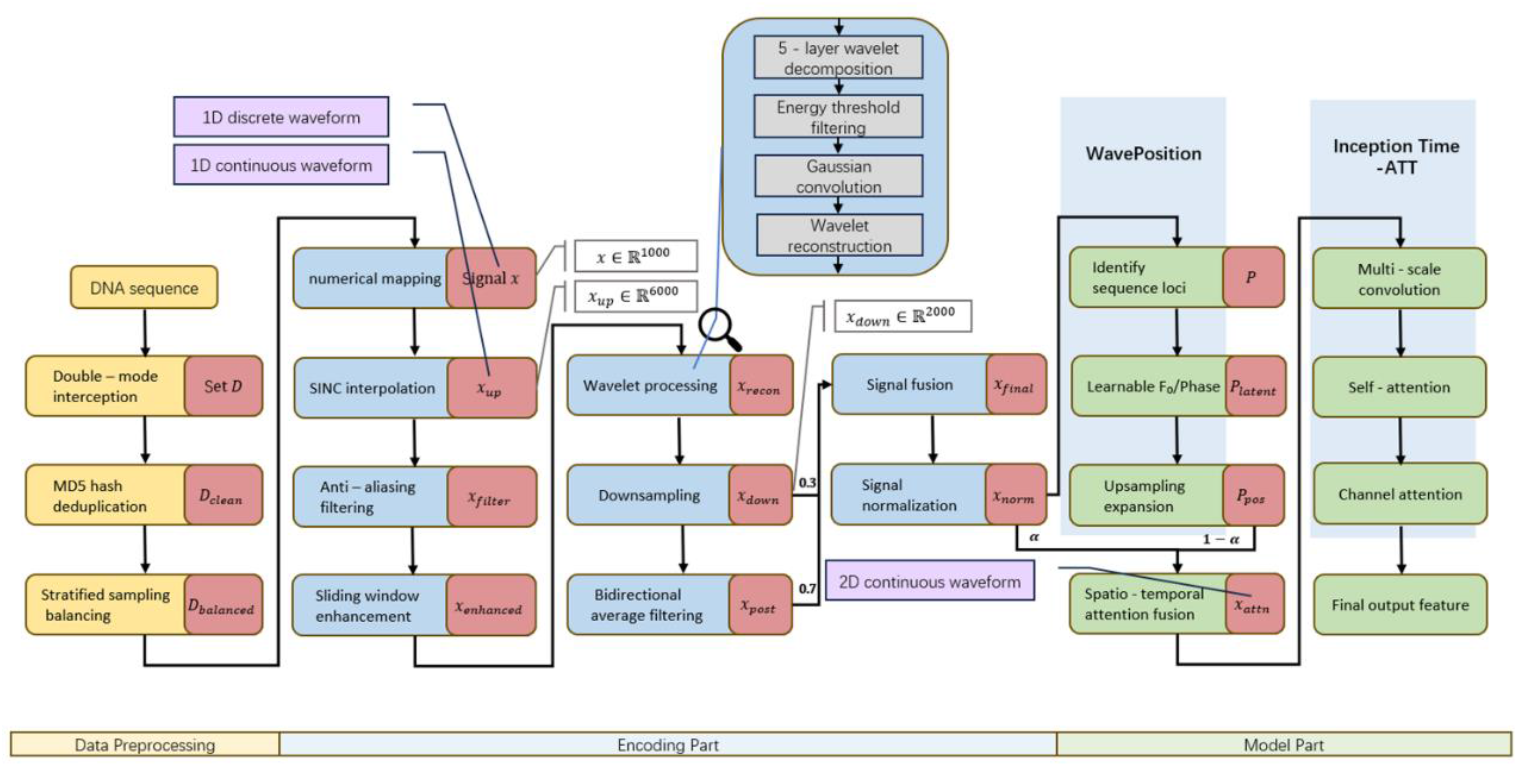
Schematic overview of the UniWave waveform encoding framework. The pipeline consists of three components: data preprocessing, the encoding framework, and the computational model. Purple boxes denote key stages where nucleotide sequences are transformed into continuous waveforms, while white boxes indicate changes in sequence length. The values 0.3, 0.7, α, and 1−α represent fusion weights. Insets illustrate the internal structure of the wavelet-based processing.

### 2.1. Data Preprocessing

### 1) Sequence Filtering and Length Normalization

Nucleotide sequences were extracted from input FASTA files, retaining only valid bases consisting of A, T, C, and G. For sequences longer than 1000 bp (*L* > 1000), denoted as *S* = {*R*_1_, *R*_2_,…, *R*_*L*_}, a random start position *s* ∈ [0, *L* − 1000] was chosen to extract a subsequence 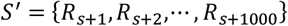. This procedure ensures coverage of potential functional domains throughout the genome (e.g., conserved regions of the NS5 non-structural protein) and reduces the risk of overfitting to fixed local fragments. The final sequence set *D* = {*d*_*i*_|*d*_*i*_ ∈ {*A, T, C, G*}^1000^, i = 1,2, …, *N*} contains strictly 1000 *bp* fragments.

#### 2) Global De-duplication and Class Balancing

A unique identifier was generated for each sequence using the MD5 hash function, *h*(*d*) = *MD*5(*d*). To eliminate redundant sequences across different categories[16, 17], a hash set was used, resulting in a non-redundant benchmark dataset, *D*_*clean*_ = {*D*_1_, *D*_2_, …, *D*_*C*_}, where *D*_*c*_ denotes the set of sequences belonging to class *c*. To address class imbalance and reduce sampling bias, stratified random sampling was performed. Taking the smallest class as the reference, an equal number of sequences were randomly selected from each class to construct a balanced dataset, denoted as *D*_*balanced*_.

### 2.2. Construction of the Continuous Waveform Encoding Framework

#### 1) Initial Mapping from Nucleotide Sequences to Waveforms

In the first stage of the UniWave encoding process, a numerical mapping function *ϕ*: {*A, T, C, G*} → ℝ is defined to convert the discrete nucleotide sequence into a discrete waveform signal. The mapping is specified as *ϕ*(*A*) = 0.1, *ϕ*(*T*) = 0.3, *ϕ*(*C*) = 0.7, and *ϕ*(*G*) = 0.9. The core principle of this mapping design is to distribute the numerical values evenly along the Y-axis, with 0.5 as the baseline (Fig. 2). Waveforms closer to the baseline (0.5) carry weaker effective information, as the signal intensity is determined by the deviation from the baseline. This naturally attenuates the influence of uncertain nucleotides (e.g., N) and inevitable noise encountered during deep learning. Regarding the numerical assignment for A, T, C, and G, we validated the scheme through controlled experiments (see Supplementary Table S7). The results indicate that when the mapping rules achieve a clear separation between the AT and CG groups, the model exhibits significantly improved classification performance. This observation suggests that the model can partially leverage the “GC bias,” a key sequence statistical feature, during learning. GC and AT differ in their intrinsic chemical structures (GC forms three hydrogen bonds, whereas AT forms two), and their distribution in the genome is often closely associated with functional region localization, fragmentation patterns, and other biological characteristics. Therefore, when numerical encoding reflects the inherent physicochemical differences of the bases, the model can more efficiently learn the mapping between sequence structure and biological function. For a DNA sequence of length 1000 *bp*, denoted *d* = {*b*_1_, *b*_2_, …, *b*_1000_}, the mapped signal is represented as *x* ∈ ℝ^1000^, where each signal value is computed as *x*_*i*_ = *ϕ*(*b*_*i*_), for *i* = 1,2, …,1000.

**Figure 2.**
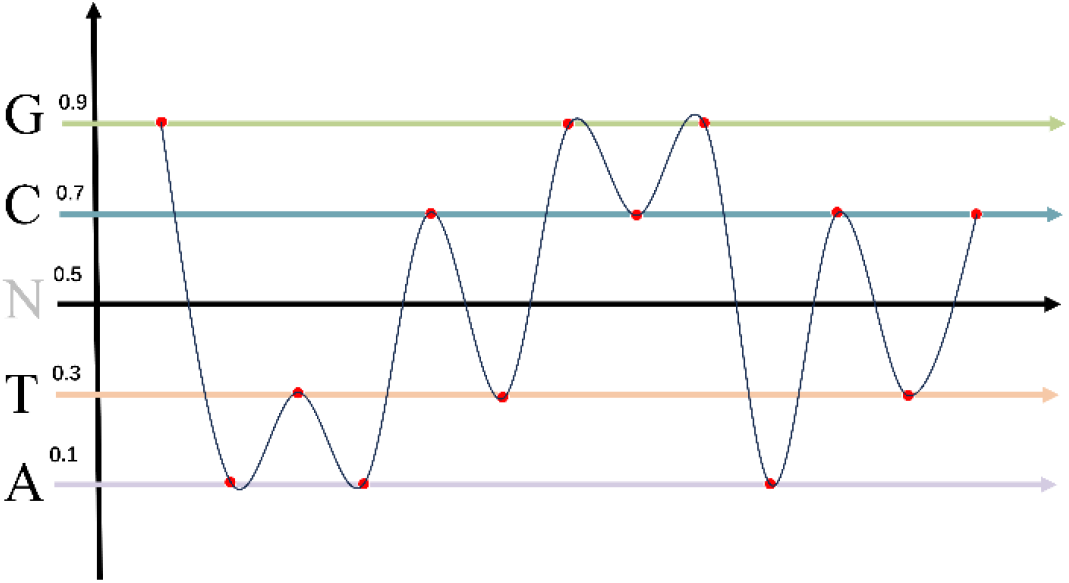
Schematic Illustration of Numerical Mapping.

#### 2) Waveform Continuization

In the second stage of UniWave encoding, the discrete nucleotide signal is reconstructed into a continuous waveform using windowed SINC interpolation, enabling alignment with the intrinsic frequency characteristics of biomolecular structures while suppressing high-frequency distortion. This process is grounded in the extended form of the Nyquist–Shannon sampling theorem, and its mathematical formulation is as follows:

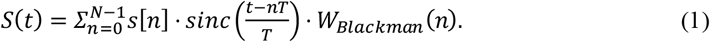

The standard SINC function, defined as 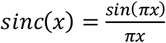, reconstructs a continuous signal through the weighted superposition of discrete sampling points. The oversampling period, 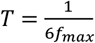, is determined by the base pair spacing of 3.4 Å in the DNA double helix, corresponding to a maximum spatial frequency of 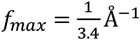. Here, s[n] denotes the discrete nucleotide signal. A 48th-order Blackman window, defined as 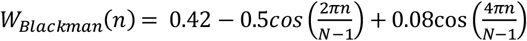, is applied to suppress high-frequency sidelobes of the interpolation kernel (Fig. 3c), reducing Gibbs oscillations to below −60 dB and markedly improving waveform smoothness.

**Figure 3.**
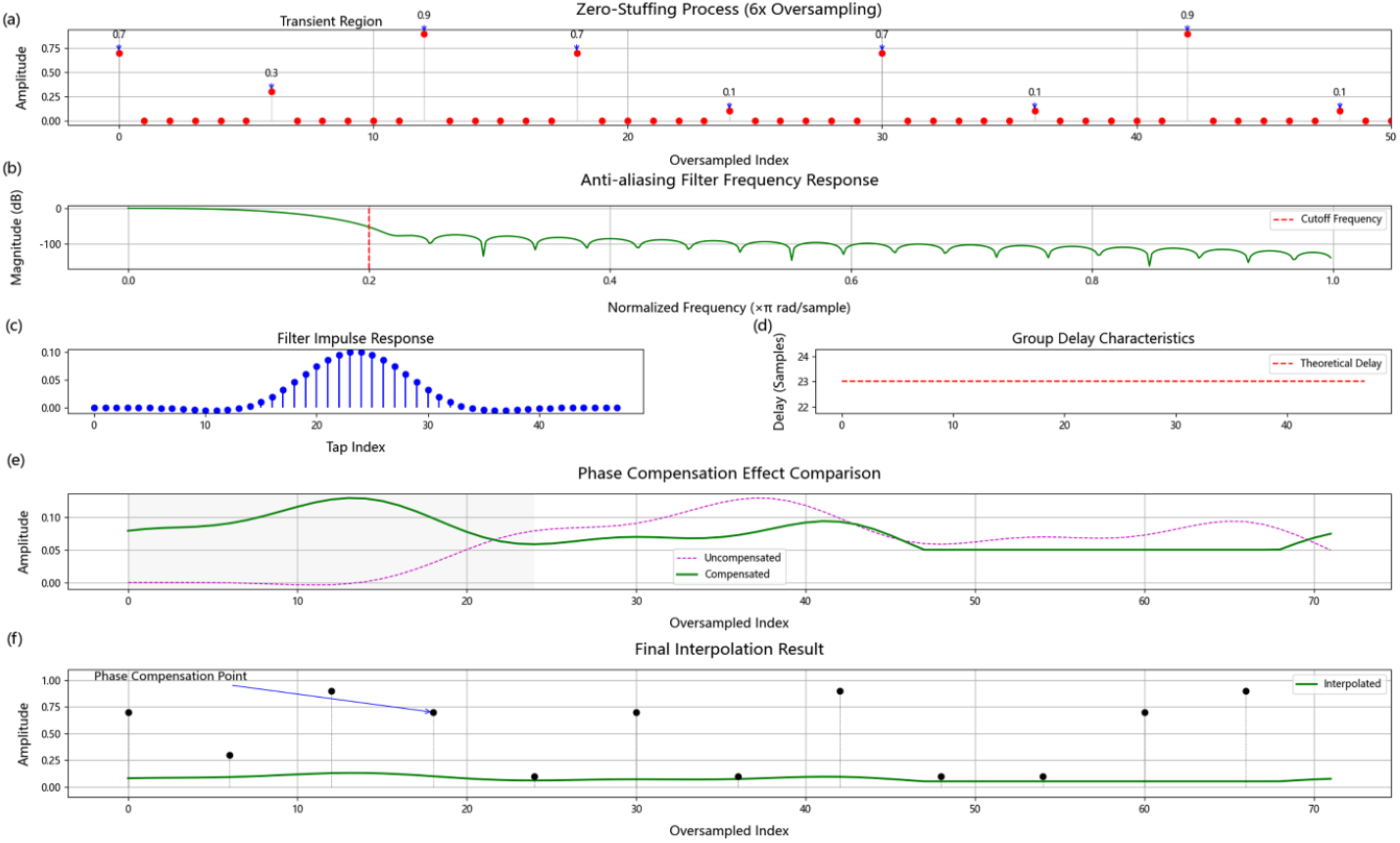
Non-equidistant SINC interpolation with anti-aliasing filtering. (a) Zero-padded signal with original sample points (red spheres). (b–d) A 48th-order Blackman-windowed FIR filter with a cutoff frequency of 0.1π rad/sample, symmetric impulse response, and 23-sample group delay compensation. (e) Comparison of filter outputs before and after delay compensation. (f) Final interpolated waveform passing exactly through the original sample points, with compensation locations indicated by blue arrows. These results visually demonstrate the effectiveness of anti-aliasing filtering and phase correction for accurate signal reconstruction.

In practice, to circumvent the high computational complexity (*O*(*N*^2^)) associated with explicitly computing the weighted SINC summation, an equivalent strategy combining oversampling and anti-aliasing filtering [18, 19] is employed to ensure signal continuity. First, the original signal *x* ∈ ℝ^1000^ is uniformly oversampled by a factor of 6 to generate a high-resolution discrete signal *x*_*up*_ = ℝ^6000^ following the procedure below:

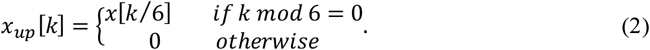

This operation redistributes the energy of the sparsely sampled original signal uniformly across higher frequency bands in the time domain (Fig. 3a), but it also generates high-frequency mirror components, commonly known as aliasing. Aliasing arises when the sampling rate is lower than the signal’s maximum frequency, causing high-frequency components to fold into lower frequencies and distort the signal. To remove these aliasing artifacts, an anti-aliasing filter is applied.

The filter design is carefully matched to the spatial frequency characteristics of DNA. The cutoff frequency is set to a normalized value of 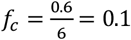, determined by the oversampling factor *K* = 6 and the maximum signal frequency *f*_*max*_. This ensures that the effective passband [0,0.1] covers the hydrogen bond vibration range between bases (0.176 Å^−1^ 0.1 × 6*f*_*max*_). The coefficients of the 48-tap Blackman-windowed filter (Fig. 3b) are obtained by truncating the ideal sinc impulse response: 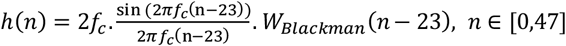, where the symmetric index shift *n* − 23 guarantees linear-phase behavior of the filter.

The filtering operation is carried out via convolution, with the filtered signal expressed as *x*_*filter*_ = *x*_*up*_ * *h*(*n*). The finite impulse response (FIR) filter introduces a group delay of *τ* = (*N* − 1)/2 = 23 samples (Fig. 3d). Phase compensation is achieved through circular shifting:

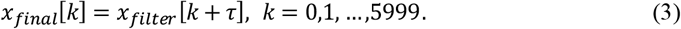

The final output is a smoothed continuous waveform *x*_*final*_ ∈ ℝ^6000^, with high-frequency components fully preserved and free from spectral aliasing (Fig. 3e–f).

To further accentuate high-variability regions (*σ*_*w*_[*i*] > 0.1) [20], such as those containing functional motifs, while moderately amplifying stable regions (Fig. 4), we developed a waveform-adaptive enhancement and local feature amplification scheme. This method suppresses noise while highlighting biologically relevant signals, thereby enhancing the intrinsic robustness of the encoded data [21, 22].

**Figure 4.**
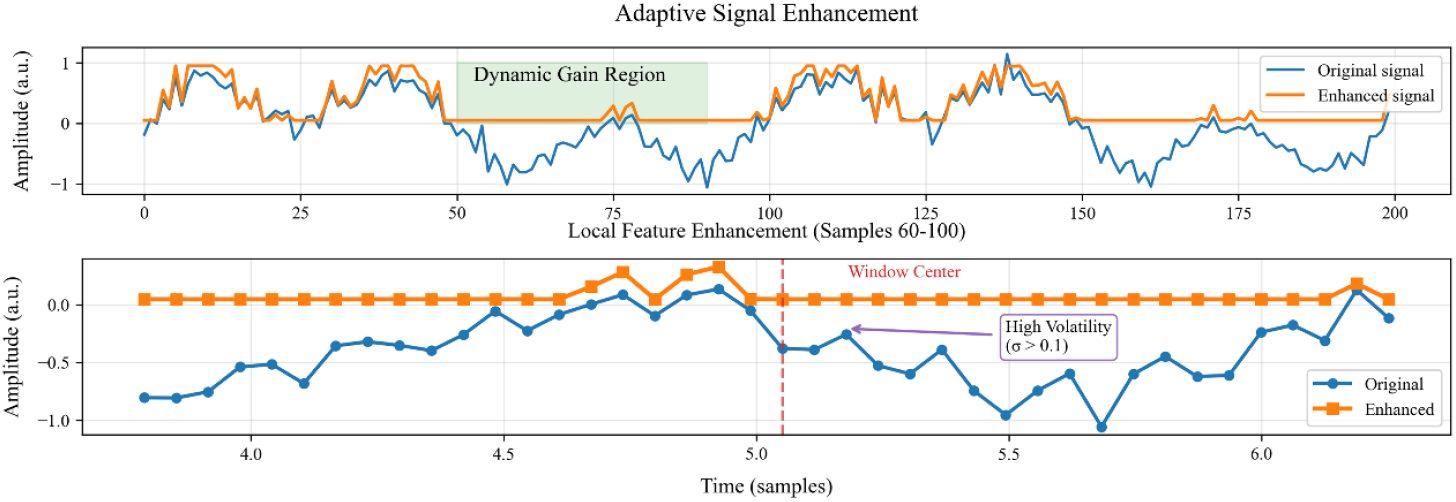
Visualization of adaptive signal enhancement. A dual-view layout illustrates enhancement effects using composite multi-frequency sine waves with added Gaussian noise. The upper panel shows the global signal (samples 0 –200), while the lower panel highlights local dynamics in a critical region (samples 60–100). Compared with the original signal (blue), the enhanced signal (orange) markedly suppresses high-frequency noise within the dynamic gain region (samples 50–90, green shading). A zoomed-in view of samples 4.0–6.0 (corresponding to global samples 60–100) marks the window center (red dashed line) and regions of high fluctuation (σ > 0.1, purple outline), where the enhanced signal exhibits smoother transitions. These multi-scale results demonstrate the algorithm’s ability to preserve global structure while optimizing local dynamics.

Local statistics were computed using a sliding window of size *w* = 60. The choice of a 60-base window is biologically motivated: 10 bases approximately correspond to 3 –4 codons (each codon comprising 3 nucleotides), representing a typical local region size in genomic sequence analysis that can capture biologically meaningful local patterns. For each position *i*, the local mean *μ*_*w*_[*i*] and standard deviation *σ*_*w*_[*i*] were calculated as:

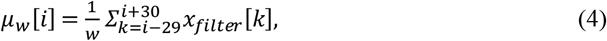

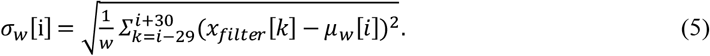

The dynamic enhancement rule is defined as:

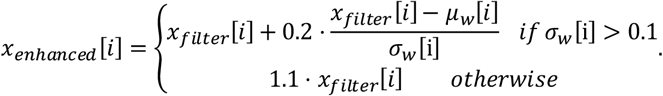

#### 3) Further Downsampling and Smoothing of the Waveform

In the third stage of the UniWave encoding process, to mitigate the substantial increase in sequence length induced by oversampling—which leads to excessive computational resource demands—further downsampling [23, 24] of the waveform is performed. To avoid significant loss of original sequence features from aggressive downsampling, we employ a wavelet transform to perform multiscale decomposition of both global trends and local features in the oversampled signal, followed by signal reconstruction. This strategy effectively preserves critical features while reducing sequence length. Furthermore, the separation and reconstruction of essential information allow for secondary extraction of the biochemical properties and spatial structural features inherent in the biological sequence. The detailed implementation steps are outlined as follows:

In this study, the enhanced signal *x*_*enhanced*_ ∈ ℝ^6000^ was decomposed into five levels using the sym5 wavelet (Fig. 5). The discrete wavelet transform (DWT), which underlies this decomposition, employs a pair of low-pass and high-pass filters, denoted as *h*[*n*] and *g*[*n*]. For a discrete signal *x*[*n*] of length *N*, decomposition at level *j* is carried out using the following operations:

**Figure 5.**
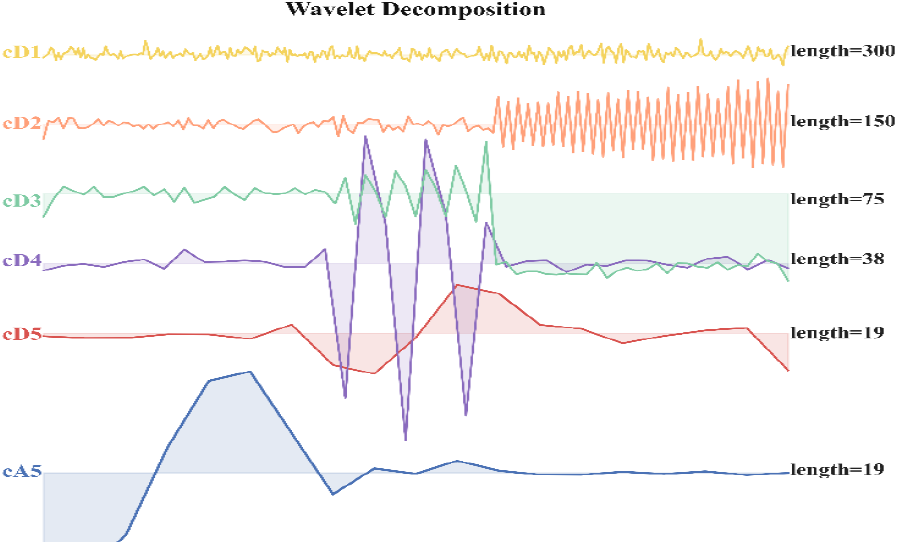
Symlet-5 wavelet multi-resolution decomposition. A five-level decomposition of a synthetic non-stationary signal composed of time-varying multi-frequency sine waves with Gaussian noise. Color gradients (dark blue to golden yellow) illustrate layered time–frequency components: the approximation cA5 (0–0.78 Hz) preserves the low-frequency signal backbone, while detail components cD5–cD1 capture transient high-frequency jumps (cD5), intermediate oscillatory patterns (cD4–cD3), and slow-varying noise (cD2–cD1). The annotated component lengths (600 → 300 → 19 points) reflect the progressive down-sampling inherent to wavelet decomposition.

Low-pass filtering followed by downsampling produces the approximation coefficients *cA*_*j*_, which capture the low-frequency components of the signal at scale *j* and reflect its overall trend. These coefficients are obtained by convolving the signal with the low-pass filter *h*[*n*] and then performing downsampling, as expressed by:

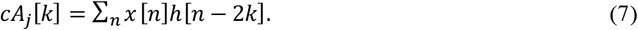

Here, 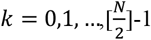; for *j* > 1, *x*[*n*] = *cA*_*j*−1_ [*n*].

High-pass filtering followed by downsampling produces the detail coefficients *cD*_*j*_, which capture the high-frequency components of the signal at scale *j*, representing local variations. These coefficients are obtained by convolving the signal with the high-pass filter *g*[*n*] and then performing downsampling, as given by:

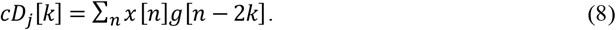

Similarly, 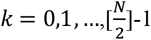. For *j* = 1, the input signal is *x*[*n*] = *x*_*enhanced*_ [*n*]; for *j* > 1, the input is the approximation coefficients from the preceding level, i.e., *x*[*n*] = *cA*_*j*−1_[*n*]. This decomposition process proceeds iteratively until the fifth level, producing the approximation coefficients *cA*_5_ and the detail coefficients *cD*_5_ (the full decomposition procedure is provided in the supplementary file titled “Wavelet Decomposition”). Ultimately, a set of coefficients is obtained, comprising *cA*_5_ and the detail coefficients *cD*_*j*_(*j* = 1,2,3,4,5), satisfying:

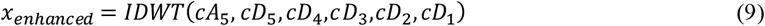

Here, IDWT denotes the inverse discrete wavelet transform, with its reconstruction formula defined using the low-pass reconstruction filter *h*_*rec*_[*n*] and the high-pass reconstruction filter *g*_*rec*_[*n*]:

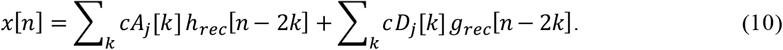

After obtaining the detail coefficients *cD*_*j*_ at each level, we applied energy thresholding to eliminate noise interference. For each set of detail coefficients *cD*_*j*_, the 95th percentile of the absolute values was computed and used as the energy threshold *τ*_*j*_:

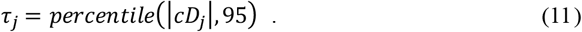

The threshold is selected based on the assumption that noise typically exhibits lower energy. Coefficients with absolute values greater than the threshold are retained using the indicator function *I*(·):

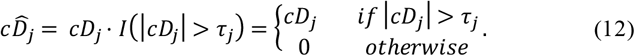

In nucleic acid sequence analysis, functional motifs are frequently associated with regions of high-frequency energy. This operation effectively preserves functionally relevant high-frequency features while removing low-energy noise arising from random sequence fluctuations.

To further smooth high-frequency details, the filtered detail coefficients 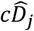 are convolved with a five-point Gaussian kernel. The kernel *K* = [0.05,0.25,0.4,0.25,0.05] satisfies the normalization condition ∑ *K* = 1. The convolution operation is defined as:

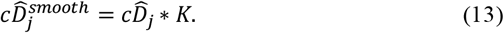

In practical implementation, zero-padding with symmetric extension is applied at the signal boundaries to preserve the original signal length.

The smoothed detail coefficients 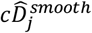 together with the original approximation coefficients *cA*_5_ are input into the inverse discrete wavelet transform (IDWT) for signal reconstruction (Fig. 6a–b), yielding a 6000-point reconstructed signal 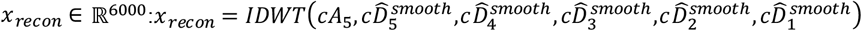

**Figure 6.**
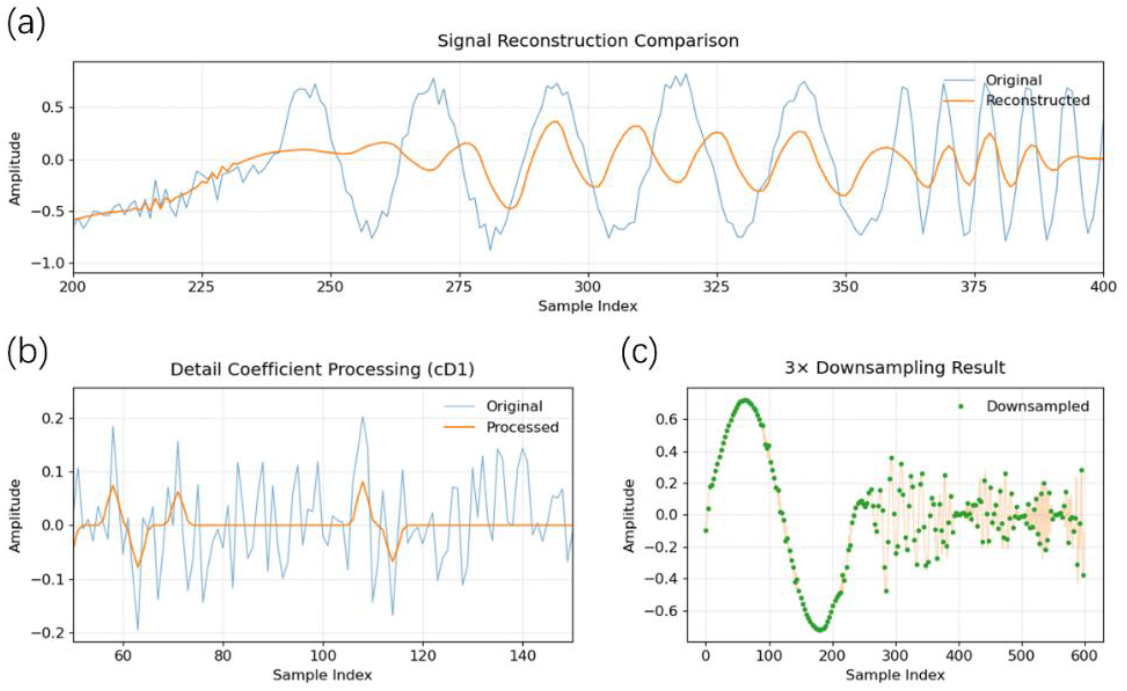
Wavelet threshold reconstruction and downsampling. The pipeline is demonstrated on a synthetic signal composed of time-varying multi-frequency sinusoidswith Gaussian noise. (a) The reconstructed signal (orange) retains key features, including a transient 5 Hz jump (t 2–3 s) and high-frequency 15 Hz oscillations (t > 3 s), while substantially suppressing noise (≈4.2 dB SNR improvement) over the 5 s duration. (b) cD1 detail coefficients after 95% thresholding and convolutional smoothing isolate informative components from background noise. (c) Three-fold downsampling reduces the signal from 600 to 200 points while preserving essential waveform characteristics, confirming effective signal compressibility.

Finally, a three-fold downsampling operation is applied to convert the 6000-point signal into a 2000-point encoded representation *x*_*down*_ ∈ ℝ^2000^ (Fig. 6c), according to:

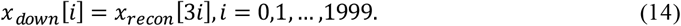

The Wavelet–Convolution Joint Compression (WCJC) method decomposes the enhanced signal into multiscale approximation and detail coefficients using wavelet transformation, suppresses noise through energy-based thresholding, and further smooths high-frequency components via Gaussian convolution. A final downsampling step is applied to optimize the dimensionality of the encoded representation. In the context of nucleic acid sequence analysis, this strategy effectively captures functionally relevant features across multiple scales, yielding a compact yet information-rich representation that serves as high-quality input for downstream modeling and analysis.

#### 4) Post-processing Filtering and Normalization of the Waveform

A bidirectional five-point moving average filter is applied to the 2000-point signal *x*_*down*_ ∈ ℝ^2000^ obtained from WCJC, using a symmetric window to suppress residual noise while preserving local edge features. The filtering operation is defined as:

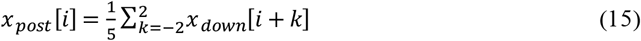

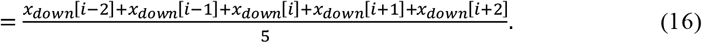

The smoothed signal *x*_*post*_ is subsequently fused with the original downsampled signal *x*_*down*_ via weighted integration to balance noise suppression and feature preservation. The final encoded representation is obtained as:

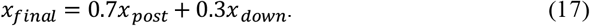

In this weighting scheme, a weight of 0.7 is assigned to the smoothed signal to suppress residual high-frequency noise that may persist following downsampling. A weight of 0.3 is assigned to the original signal to preserve local detail features, preventing excessive smoothing and ensuring retention of critical information.

To finalize signal preprocessing prior to model input, the fused signal *x*_*final*_ is normalized as follows:

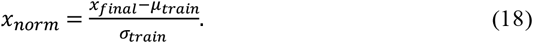

### 2.3. Construction of a Attention-Enhanced Deep Learning Model 1)Learnable Positional Encoding Module (WavePosition)

To prevent potential loss of positional information in nucleic acid sequences during the UniWave encoding process, we developed a dedicated positional encoding module, termed WavePosition. Within the spatiotemporal attention–enhanced deep model, the learnable WavePosition module embeds sequence positional information into continuous waveform signals, overcoming the limitation of conventional models that are insensitive to absolute positional context. The module generates positional features via learnable frequency–phase parameters and dynamically fuses these features with sequence signals through a spatiotemporal attention mechanism. he detailed implementation is described as follows:

Define the learnable base frequency parameter *w* ∈ ℝ^1×1×*L*^ and the phase parameter *Φ*_0_ ∈ ℝ^1×1×*L*^, where *L* = 20 denotes the latent dimension used to generate the initial positional features. For each position *t* ∈ [1, *T*] in the sequence (with *T* = 2000 representing the signal length), the initial positional features are generated using a sine function:

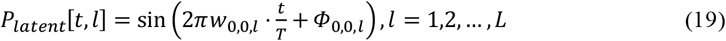

Here, *w*_0,0,*l*_ denotes the base frequency of the *l*-th latent dimension, controlling the oscillation frequency of the positional signal, which reflects the periodic distribution of functional regions in nucleic acid sequences, such as the positional preference of promoter regions. *Φ*_0,0,*l*_ represents the initial phase, introducing position-specific phase shifts, for example, capturing local fluctuations at splice sites.

By initializing these parameters with uniform and normal distributions, the model is enabled to capture and explore a diverse range of positional patterns.

As the initial positional feature has a dimensionality of *L* = 20, it must be expanded to match the sequence length *T* = 2000 using linear interpolation. The upsampling scale factor is defined as 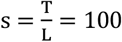, and a linear interpolation function *Upsample*(·, *s*) is applied to generate the final positional encoding *P*_*pos*_ ∈ ℝ^*T*^, with *P*_*pos*_[*t*] = *Upsample*(*P*_*latent*_, *s*)[*t*]. The normalized signal *x*_*norm*_ ∈ ℝ^*T*^ is concatenated with the positional encoding *P*_*pos*_ ∈ ℝ^*T*^ along the channel dimension, forming the input feature matrix *X* = [*x*_*norm*_, *P*_*pos*_] ∈ ℝ^2×*T*^. A dual-layer convolutional network is subsequently used to compute attention weights *α* ∈ ℝ^*T*^, facilitating the adaptive fusion of the signal and positional information.

WavePosition establishes a position-sensitive modeling framework using learnable frequency–phase parameters, addressing the limitation of conventional convolutional neural networks in capturing absolute positional information. The spatiotemporal attention mechanism dynamically adjusts weights based on signal variability, preserving detailed information in highly variable regions to enhance mutation detection, while emphasizing positional priors in conserved regions to capture functional motif specificity. By compressing the latent positional dimension to *L* = 20, the model maintain spositional information while reducing the parameter scale of the positional encoding by two orders of magnitude. Differentiable upsampling then expands the compressed encoding to the full sequence length, avoiding exponential growth of computational complexity in high-dimensional space and providing a topology-aware embedding that enables efficient multi-scale feature extraction within the downstream InceptionTime-ATT module.

#### 2)Hybrid Attention Network (InceptionTime-ATT)

As the primary feature extraction module, the hybrid attention network integrates multi-scale convolutions [25, 26], self-attention [27, 28], and channel attention [29], enabling hierarchical capture of local sequence motifs, long-range dependencies, and inter-channel feature relevance in nucleic acid sequences. Moreover, it employs lightweight operations, including depthwise separable convolutions and low-dimensional attention computations, to optimally balance representational power with computational efficiency.

## 3. Results and discussion

### 3.1. Experimental Setup and Evaluation Metrics

#### 1)Datasets and Experimental Grouping

We evaluated the UniWave framework using four genomics datasets with clear biological relevance, enabling a systematic assessment of performance, robustness, versatility, and generalization. Each dataset was selected to address a distinct evaluation objective, ensuring comprehensive validation. Key dataset statistics are summarized in Table I.

**Table 1.**
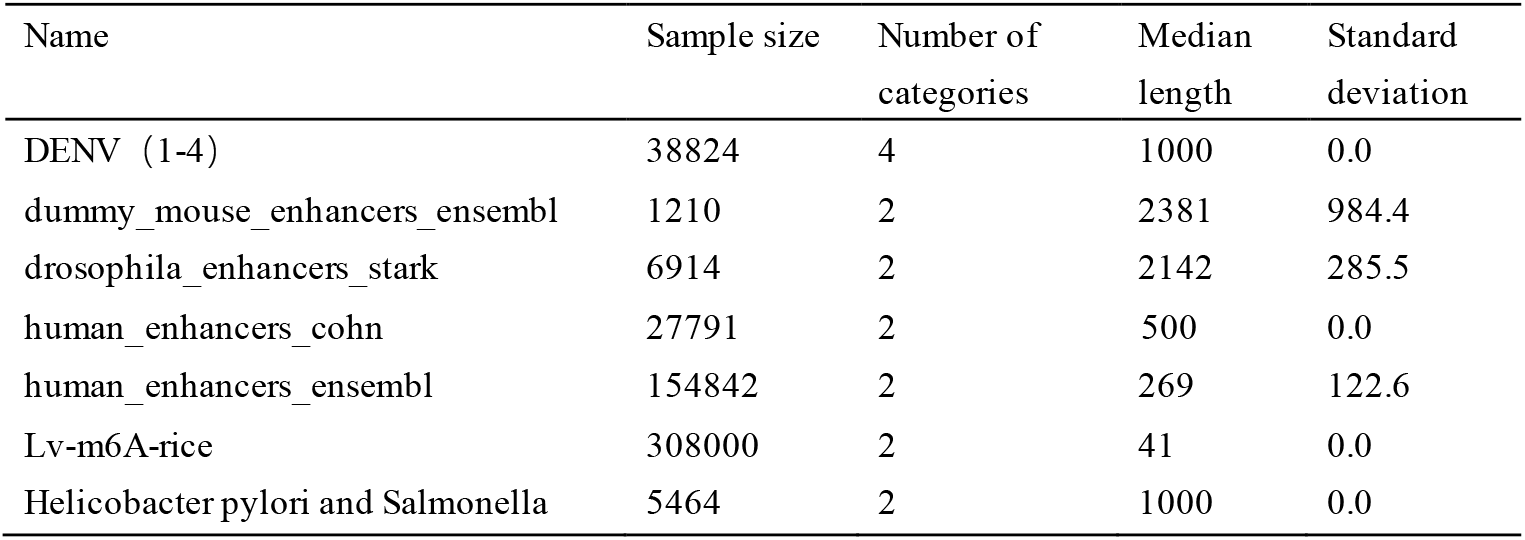
Key dataset statistics.

| Name | Sample size | Number of categories | Median length | Standard deviation |
| --- | --- | --- | --- | --- |
| DENV ( 1-4 ) | 38824 | 4 | 1000 | 0.0 |
| dummy_mouse_enhancers_ensembl | 1210 | 2 | 2381 | 984.4 |
| drosophila_enhancers_stark | 6914 | 2 | 2142 | 285.5 |
| human_enhancers_cohn | 27791 | 2 | 500 | 0.0 |
| human_enhancers_ensembl | 154842 | 2 | 269 | 122.6 |
| Lv-m6A-rice | 308000 | 2 | 41 | 0.0 |
| Helicobacter pylori and Salmonella | 5464 | 2 | 1000 | 0.0 |

The Dengue virus (DENV) genome dataset was obtained from NCBI and includes serotypes 1–4 (classes 0–3). A strictly balanced subset was constructed to eliminate class bias, focusing on the envelope (E) protein coding region, which is closely linked to viral antigenicity and evolutionary divergence. As a canonical target for viral genotyping, this region provides a rigorous benchmark for evaluating model robustness and noise tolerance under complex perturbations.

To further validate the framework across tasks and species, three additional datasets were incorporated. First, a cross-species enhancer classification dataset from the Genomic Benchmarks repository (BMC Genomics Data[30]) was used, comprising four subsets spanning mouse, Drosophila, and human genomes. The high sequence heterogeneity and broad species coverage closely reflect real-world genomic variability and enable evaluation of cross-species adaptability. Second, the rice m6A benchmark dataset (Lv-m6A-rice[31]), consisting of 41-bp sequences with a fixed central adenine, was employed to assess performance on plant epigenetic modification detection. Its widespread use allows direct comparison with established tools, validating the applicability of waveform encoding to epigenetic site identification. Third, a bacterial classification dataset comprising Helicobacter pylori and Salmonella genomes from NCBI was constructed to examine cross-domain generalization from viral to bacterial genomes. To analyze the hierarchical contributions of the waveform encoding framework, three comparative encoding variants were designed using the DENV dataset (Fig. 7). The baseline model maps A/T/C/G to 0.1/0.3/0.7/0.9 based on hydrogen bond properties, yielding a 1×1000 discrete feature vector with biological interpretability. The intermediate model applies 6× SINC interpolation with anti-aliasing filtering to generate a 1×6000 continuous waveform, enhancing smoothness and reducing aliasing. The full model further incorporates adaptive signal enhancement and a wavelet–convolution joint compression (WCJC) module, compressing the representation to 1×2000 dimensions while preserving salient biological features.

**Fig. 7.**
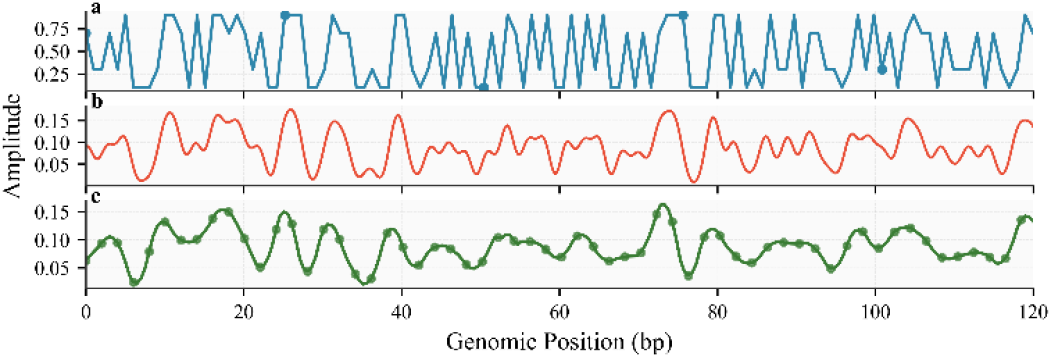
Complete genomic encoding and processing workflow. (a) Raw encoding (blue) maps A/T/C/G bases to fixed values in the 0–120 bp region (0.1, 0.3, 0.7, 0.9). (b) SINC interpolation (red; 6× oversampling) increases waveform smoothness by 79%. (c) Wavelet–convolution compression (green; 3× downsampling) reduces data volume by 66.7% while preserving key feature peaks, highlighting the effectiveness of time–frequency processing for genomic representation.

All variants share the same backbone architecture, including the learnable WavePosition module and the InceptionTime-ATT network.

#### 2) Anti-Interference Validation Framework

We implemented a two-stage data augmentation strategy to rigorously evaluate model robustness (Supplementary Table 1). Mild perturbations were introduced to the training set (Supplementary Fig. 1), whereas the challenge test set was exposed to amplified noise (Supplementary Fig. 2) to assess generalization under extreme conditions. The effectiveness of this augmentation strategy was further assessed using multiple metrics, including global and local waveform comparisons (Supplementary Fig. 3a) and analyses of data distribution expansion (Supplementary Fig. 3b–c).

#### 3) Evaluation Metrics

To meet the biological sensitivity demands inherent to viral genotyping tasks, we selected three evaluation metrics:

a. Balanced Accuracy (BACC)

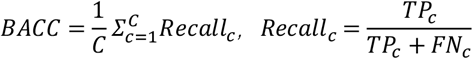

Balanced Accuracy (BACC) computes the average recall across classes, mitigating class imbalance and reflecting the model’s ability to recognize all classes fairly.
b. Macro-averaged F1 Score (F1-macro)

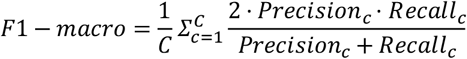

The F1 score balances precision and recall and is especially sensitive to underrepresented classes, such as rare mutant sequences.
c. Macro-averaged Multi-class AUC-ROC Using a One-vs-Rest scheme, the AUC-ROC was computed for each class and then macro-averaged across all classes: 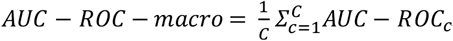.

### 3.2. Evaluation of UniWave’s Robustness to Noise

We systematically evaluated UniWave’s robustness to noise along three dimensions: intrinsic noise suppression, robustness under augmented training, and performance relative to conventional encoding methods. All experiments focused on the challenge test set (specific augmentation parameters are provided in the Supplementary Materials) to highlight core evaluation objectives. Under a no-augmentation training protocol (Table II), the baseline and intermediate models achieved balanced accuracies (BACC) of 66.31 % and 66.16%, respectively, with no significant difference, indicating that interpolation-based anti-aliasing alone is insufficient to substantially enhance intrinsic noise resilience. In contrast, the full model—integrating adaptive signal enhancement, local feature amplification, and the WCJC downsampling module—achieved a BACC of 72.98% through its multi-scale waveform analysis architecture, demonstrating the advantage of targeted encoding-layer optimization in complex noisy environments. The full model contains only 27.2K parameters with a 1×2000 feature dimension, outperforming the intermediate model in both parameter efficiency and feature compactness, underscoring the effectiveness of the lightweight design.

**Table 2.** Noise resistance of encoding strategies without training augmentation.

| Encoding Stage | Model Params (K) | Feature Dimension | Challenge Test (BACC) |
| --- | --- | --- | --- |
| Baseline Model | 25.6 | $1 \times 1000$ | 66.31% (95% CI: 0.6419-0.6845) |
| Mid Model | 33.9 | 1×6000 | 66.16% (95% CI: 0.6426-0.6811) |
| Full Model | 27.2 | 1×2000 | 72.98% (95% CI: 0.7048-0.7522) |

Under the two-stage augmentation framework (four standard augmentations during training; three extreme perturbations during testing, Table III), robustness was further validated. The baseline model’s BACC rose to 83.55%, yet misclassifications persisted in highly variable classes. The intermediate model, combining multi-scale interpolation with structural modeling, achieved a BACC of 98.64% and a macro-average AUC-ROC of 0.9979, with a substantial reduction in errors. These results demonstrate that the synergy of architectural enhancements and data augmentation markedly improves generalization. Although the full model’s BACC (92.22%) was slightly lower than the intermediate model, it retained its lightweight advantage. All models showed reduced performance for the DENV_3 serotype, consistent with high E protein sequencevariability dueto antigenicdrift, emphasizing the importance of mutation sites in sequence encoding. Narrow 95% confidence intervals across models indicate low data dispersion and stable, reliable performance.

**Table 3.** Model robustness under a two-stage augmentation framework.

| Encoding Stage | Model Params (K) | Feature Dimension | Challenge Test (BACC) |
| --- | --- | --- | --- |
| Baseline Model | 25.6 | 1×1000 | 83.55% (95% CI: 0.8172-0.8546) |
| Mid Model | 33.9 | 1×6000 | 98.64% (95% CI: 0.9799-0.9923) |
| Full Model | 27.2 | 1×2000 | 92.22% (95% CI: 0.9082-0.9358) |

Comparative experiments with conventional encoding methods (Table IV) further highlight UniWave’s advantages. The full model (27.2K parameters, 1×2000 features) achieved a BACC of 92.22%, whereas one-hot encoding (26.9K parameters, 4×1000 features) and word embeddings (1,044.3K parameters, 512×1000 features) reached only 77.51% and 72.41%, respectively, revealing a pronounced performance gap. These findings indicate that while conventional encodings may perform well on clean datasets, their robustness deteriorates under complex noise. In contrast, UniWave’s waveform encoding balances noise resilience and computational efficiency. The lightweight full model is suitable for current small-scale datasets and promises scalability for larger datasets. Furthermore, differences between intermediate and full models provide flexible options for varying computational resources and application requirements.

**Table 4.**
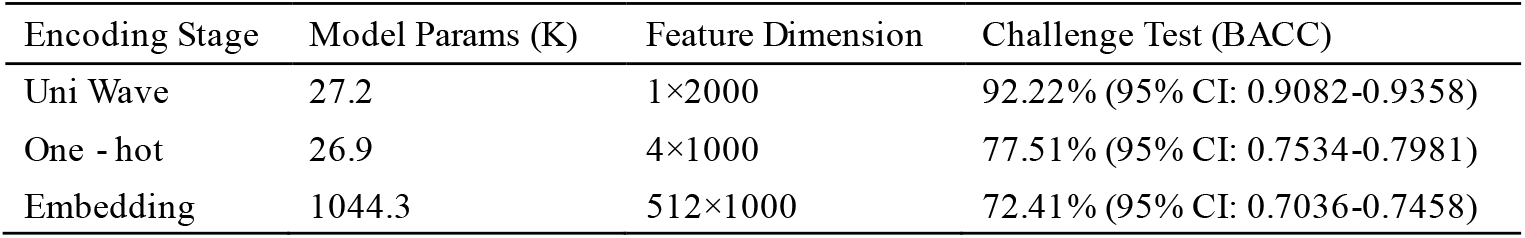
Comparison with traditional encodings under a two-stage augmentation framework.

| Encoding Stage | Model Params (K) | Feature Dimension | Challenge Test (BACC) |
| --- | --- | --- | --- |
| Uni Wave | 27.2 | 1×2000 | 92.22% (95% CI: 0.9082-0.9358) |
| One - hot | 26.9 | 4×1000 | 77.51% (95% CI: 0.7534-0.7981) |
| Embedding | 1044.3 | 512×1000 | 72.41% (95% CI: 0.7036-0.7458) |

### 3.3. Efficient Identification of Enhancer Regions by UniWave

To systematically validate the advantages of UniWave’s waveform encoding in enhancer sequence classification, we selected enhancer datasets from the BMC Genomics Data study “Genomicbenchmarks: a collection of datasets for genomic sequence classification”, providing a rigorous benchmark for encoding comparisons. Experiments used the official CNN with word embedding as the baseline and evaluated performance via accuracy (ACC) and F1 score, focusing on each encoding scheme’s adaptability to diverse enhancer sequences.

Results (Table V) show that UniWave outperforms traditional word embedding CNNs in most cases. On the dummy_mouse_enhancers_ensembl dataset, UniWave achieved an ACC of 72.73% and F1 of 72.72%, exceeding the baseline (69.0% ACC, 70.4% F1), indicating more precise capture of sequence features. The advantage was particularly pronounced on the drosophila_enhancers_stark dataset, with ACC improving from 58.6% to 68.30% and F1 from 44.5% to 67.92%. This improvement reflects the limitations of word embedding encoding in accommodating high sequence heterogeneity, whereas UniWave’s multi-scale waveform analysis effectively addresses class imbalance and feature complexity. For the human_enhancers_ensembl dataset, UniWave’s F1 reached 70.02%, markedly higher than the baseline (56.5%), confirming its superiority in capturing deep, interrelated features. Performance was comparable only on the human_enhancers_cohn dataset (UniWave ACC 67.29%, F1 67.29%; word embedding CNN ACC 69.5%, F1 67.1%), reflecting the dataset’s conserved sequences and low feature discriminability.

**Table 5.** Enhancer region identification.

| Datasets | Embedding CNN |  | Uniwave |  |
| --- | --- | --- | --- | --- |
|  | ACC | F1 | ACC | F1 |
| dummy_mouse_enhancers_ensembl | 69.0 | 70.4 | 72.73 | 72.72 |
| drosophila_enhancers_stark | 58.6 | 44.5 | 68.30 | 67.92 |
| human_enhancers_cohn | 69.5 | 67.1 | 67.29 | 67.29 |
| human_enhancers_ensembl | 68.9 | 56.5 | 70.11 | 70.02 |

Fundamentally, word embedding relies on predefined sequence fragment mappings, making it vulnerable to high genomic variability and noise, limiting generalization on complex datasets. In contrast, UniWavedynamically modelswaveform features without fixed mappings, adaptively capturing intrinsic structural patterns across species and sources. Its lightweight architecture (low parameter count, compact feature dimension) enhances feature representation while maintaining computational efficiency. Overall, waveform encoding demonstrates superior robustness and adaptability compared with traditional word embeddings, particularly for highly heterogeneous and complex datasets. UniWave retains advantages in noise-resistant scenarios and, through innovative encoding, provides an efficient and stable solution for genomic sequence classification, while offering guidance for future dataset-specific encoding optimizations.

### 3.4. Identification of m6A Sites in Rice

Using a publicly available rice m6A site identification framework, we compared UniWave with mainstream tools iDNA6mA-Rice and SNNRice6mA to assess the feasibility and effectiveness of the novel waveform encoding approach. As shown in Table VI, UniWave achieved key metrics—accuracy (ACC) and area under the curve (AUC)—that match or surpass existing tools: ACC reached 92.16%, slightly higher than iDNA6mA-Rice (91.7%) and SNNRice6mA (92.0%), while the AUC was 0.97, equivalent to SNNRice6mA and second only to the optimal benchmark. Notably, UniWave also reports a 95% confidence interval (91.94%– 92.36%), demonstrating stable and reliable predictive performance, whereas mainstream tools do not provide confidence estimates, underscoring UniWave’s experimental rigor.

**Table 6.**
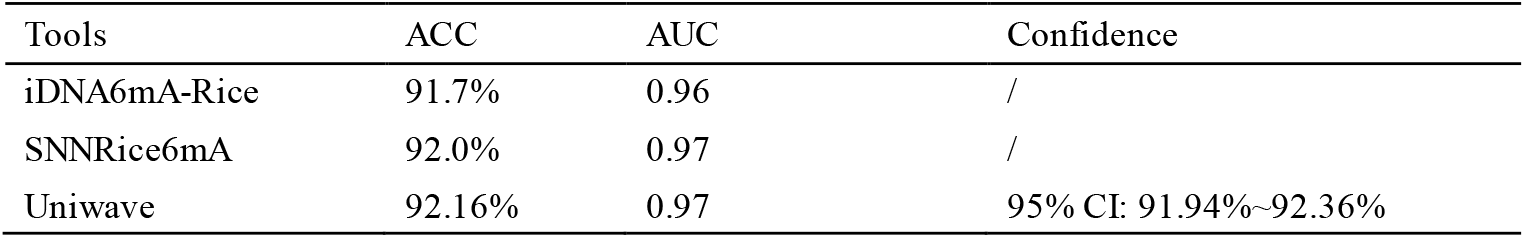
Rice m^6^A site detection.

| Tools | ACC | AUC | Confidence |
| --- | --- | --- | --- |
| iDNA6mA-Rice | 91.7% | 0.96 | / |
| SNNRice6mA | 92.0% | 0.97 | / |
| Uniwave | 92.16% | 0.97 | 95% CI: 91.94%~92.36% |

The primary significance of this experiment lies not in the marginal ACC improvement, but in validating the application of a novel encoding strategy for rice m6A site detection. While conventional tools rely on fixed sequence encodings, UniWave dynamically captures intrinsic structural features through waveform encoding without predefined mappings or complex feature engineering, achieving comparable or superior performance. The lightweight architecture (low parameter count, compact feature dimension) ensure s computational efficiency while maintaining high accuracy, offering both an innovative and practical reference for future m6A identification tools and informing the design of encoding strategies for other epigenetic modification detection tasks.

### 3.5. Contribution Analysis of Each Component in the Model

To evaluate the contributions of individual components, we performed ablation experiments on the full model under the dual-stage augmentation framework. As shown in Table 7, removing the WaveEncoder module reduced BACC by 1.23% on the standard test set and 2.02% on the challenge set, highlighting its fundamental role in model performance. In contrast, removing the channel attention module alone had a minor effect, reflecting its auxiliary role. Interestingly, removing the self-attention module alone slightly improved performance on both test sets; however, removing both self-attention and channel attention simultaneously caused a sharp 5.62% drop in BACC on the challenge set. These results underscore the synergistic importance of the two attention modules: self-attention captures long-range dependencies, while channel attention filters salient features, jointly forming a global-local interaction mechanism critical for robust feature representation.

**Table 7.**
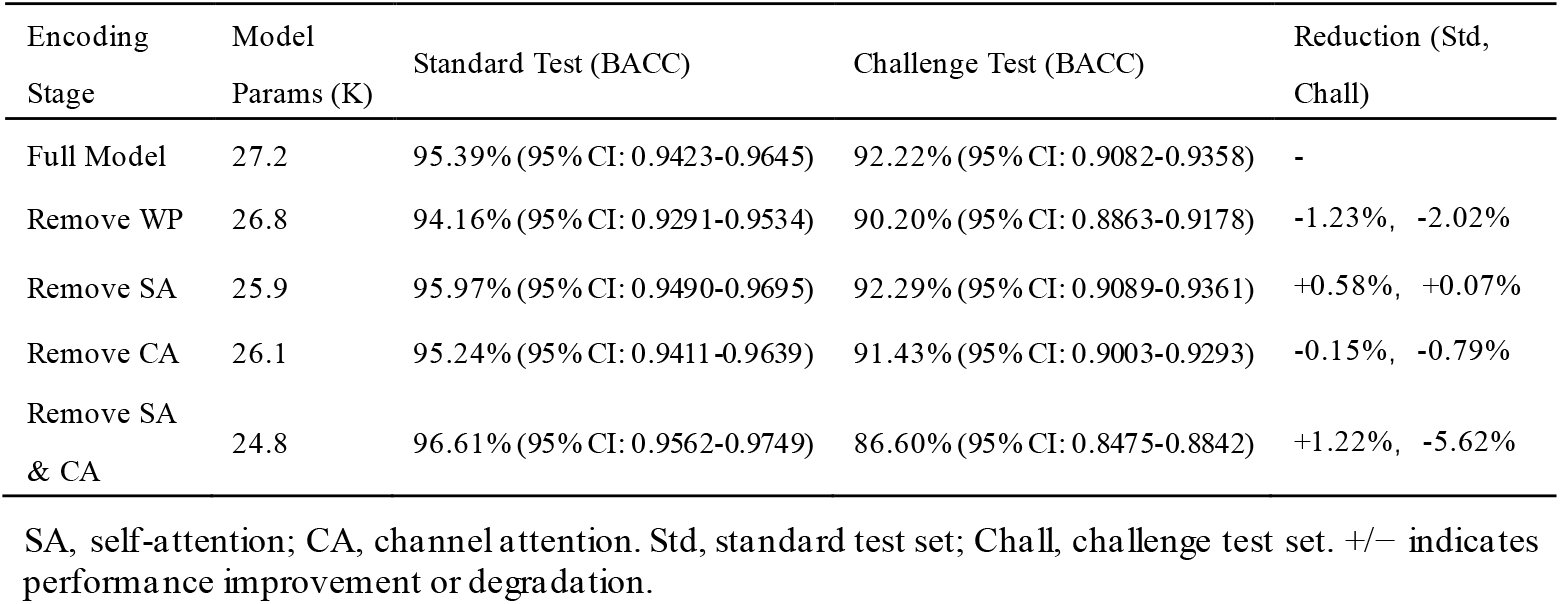
Ablation of key components in the waveform encoding architecture.

Further experiments showed that, unlike the substantial performance variations observed in the three-stage encoding experiments (Sections 3.2–3.3), ablations in model training components caused only minor fluctuations. This confirms the pivotal role of waveform-based encoding in enhancing intrinsic noise resistance. Overall, the ablation study quantifies each module’s independent contribution and reveals the synergy between waveform encoding and attention mechanisms, providing a clear roadmap for prioritizing future model optimization.

### 3.6. Cross-Species Generalization Test

To further evaluate the cross-species generalizability of waveform encoding, we constructed a bacterial classification task using Helicobacter pylori and Salmonella genomic datasets. Retaining the dual-stage augmentation protocol from the viral benchmark, we assessed the model’s ability to transfer across domains from viral to bacterial genomes.

As shown in Table 8, the full model achieved a BACC of 93.61% on the standard test set and 92.34% on the challenge set, with macro F1 and AUC-ROC scores also remaining high. These results demonstrate that the waveform-based full model accurately classifies sequences across distinct species, maintains robustness under noisy conditions, and preserves a compact parameter size and feature dimension. This confirms its generalization capability across diverse genomic contexts and provides strong evidence for the applicability of waveform encoding in multi-species scenarios, highlighting its versatility and reliability in complex biological data analysis.

**Table 8.** Cross-species generalization of waveform encoding in bacterial genome classification.

| Encoding Stage | Model Params (K) | Feature Dimension | Standard Test (BACC) | Challenge Test (BACC) |
| --- | --- | --- | --- | --- |
| Full Model | 27.0 | 1×2000 | 93.61% (95% CI: 0.9158-0.9552) | 92.34% (95% CI: 0.8988-0.9452) |

## 4. Conclusions

This study addresses key challenges in nucleic acid sequence feature extraction—namely, the curse of dimensionality, insufficient modeling of long-range dependencies, and limited noise resilience—by introducing UniWave, a novel encoding framework based on dynamic waveform modeling. UniWave transforms discrete nucleotide sequences into continuous, biophysically meaningful waveforms, integrating a learnable WavePosition encoding module with a lightweight dual-attention InceptionTime-ATT model. This design preserves biological interpretability while balancing computational efficiency and model performance.

Through multidimensional validation across four core tasks—dengue virus serotype classification, cross-species enhancer identification, rice m6A site detection, and bacterial genome classification—UniWave demonstrates clear advantages. It exhibits strong noise robustness under complex perturbations, with waveform encoding significantly outperforming traditional one-hot and word embedding approaches. Across species and tasks, UniWave shows high adaptability and generalization, achieving performance comparable to or exceeding mainstream tools with only 27.2K parameters and a compact 1×N feature dimension. Ablation studies confirm that the waveform encoding module is central to intrinsic noise resistance, while the synergy of self-attention and channel attention enhances overall robustness. Feature map visualizations (e.g., Supplementary Fig. 7) further demonstrate that UniWave preserves core information while optimizing and simplifying signal representation.

The intermediate and full model variants offer flexible solutions for high-precision or resource-constrained scenarios. Future work will explore the relationships between waveform features and biological processes, such as epigenetic regulation and protein conformational dynamics, advancing the application of this physically inspired modeling approach in precision medicine, synthetic biology, and related fields. Additionally, we aim to investigate the feasibility of constructing lightweight pre-trained models based on UniWave, seeking to approach the representational power of large-scale models like DNABERT-2 while maintaining biophysical interpretability.

## Acknowledgements

This study was funded by Gansu Province Higher Education Innovation Fund of China (2024A-005).

## Code Availability

The implementation of the UniWave framework, including related code and supporting datasets, is publicly available under the MIT License at: https://github.com/qcy7226-create/Uniwave.1

