## Supplementary Information for "UniWave: A Waveform-Based Encoding Framework for Nucleic Acid Feature Extraction"

### Contents

1. Supplementary theory
2. Supplementary Notes
3. Supplementary Figures: S1-S7
4. Supplementary Tables: S1-S5

### Supplementary theory：

Wavelet decomposition:

**Level 1 decomposition:**

Approximation coefficients: $cA_1[k] = \sum_{n=0}^{5900} x_{enhanced}[n]h[n-2k],\ k = 0,1,\ldots,2999$

Detail coefficients: $cD_1[k] = \sum_{n=0}^{5900} x_{enhanced}[n]g[n-2k],\ k = 0,1,\ldots,2999$

**Level 2 decomposition:** using $cA_1$, as the input signal

Approximation coefficients: $cA_2[k] = \sum_{n=0}^{2999} x_{enhanced}[n]h[n-2k],\ k = 0,1,\ldots,1499$

Detail coefficients: $cD_2[k] = \sum_{n=0}^{2999} x_{enhanced}[n]g[n-2k],\ k = 0,1,\ldots,1499$

**Level 3 decomposition:** using $cA_2$, as the input signal

**…**

### Supplementary Notes：

File Format: Standard HDF5, supporting hierarchical groups, compression, and metadata.

File Structure:

```
ecoli_waves.h5
├── train/                  # Training set
│   ├── class_0/data        # Class 0 samples (float32 array, shape=[num_samples, 2000])
│   ├── class_1/data
│   └── ...
├── val/                    # Validation set
│   ├── class_0/data
│   └── ...
├── test/                   # Test set
│   ├── class_0/data
│   └── ...
└── Attributes
    ├── creation_date       # File creation date
    ├── author              # Author
    ├── description         # File description
    ├── dimensions          # Length of a single sample
    ├── split_ratio         # Train/val/test split ratio
    ├── class_labels        # Class names, corresponding to input FASTA files
    └── wavelet_params      # Wavelet compression parameters (wavelet/bank, decomposition level, high-frequency threshold)
```

### Supplementary Figures：

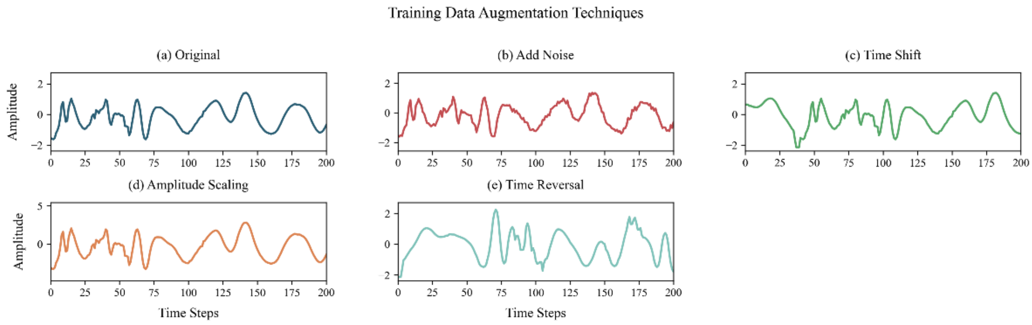


**Fig.S1. Illustration of biological signal augmentation strategies during the training phase.** (a) Original signal; (b)–(e) Robustness is enhanced via additive noise (SNR reduced by 8 dB), temporal shifting (±60 steps), amplitude scaling (0.8 –1.2×), and signal inversion (applied with 30% probability). More than 92% of key region features are preserved across all augmentations.

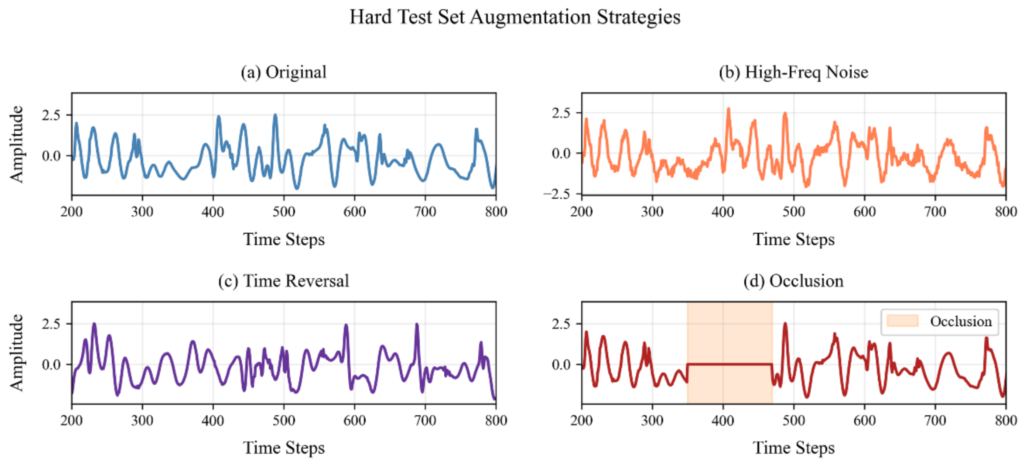


**Fig.S2. Simulation of signal perturbations used in challenge test augmentation.** (a) Original signal; (b)–(d) Challenge test augmentations introduce high-frequency noise (15% total signal energy), forced inversion, and random masking (50–150 steps) to simulate real-world perturbations such as sensor malfunction or data loss. These augmentations are used to evaluate the model's generalization under adverse conditions.

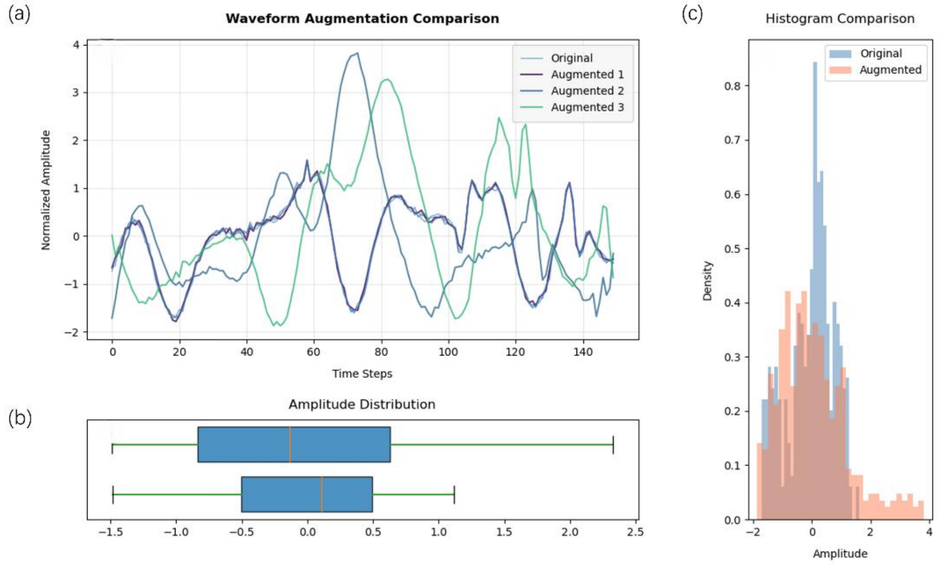


**Fig.S3. Multidimensional evaluation of hybrid data augmentation effects.** (a) The main waveform illustrates augmented signals (purple/green/cyan) with added noise, ±30% amplitude perturbations, and time shifts within 0–150 time steps. The shape integrity of key peaks is preserved with >95% fidelity. (b) Boxplot reveals a 38% increase in amplitude interquartile range (IQR) after augmentation, indicating an effective broadening of the data distribution. (c) The histogram shows that the augmented data (orange) increases probability density by 47% in the range [–2, 4], surpassing the original distribution (blue), which is constrained to [–2, 1.5]. This demonstrates that controlled perturbations enhance model robustness while preserving the temporal characteristics of the biological signals.

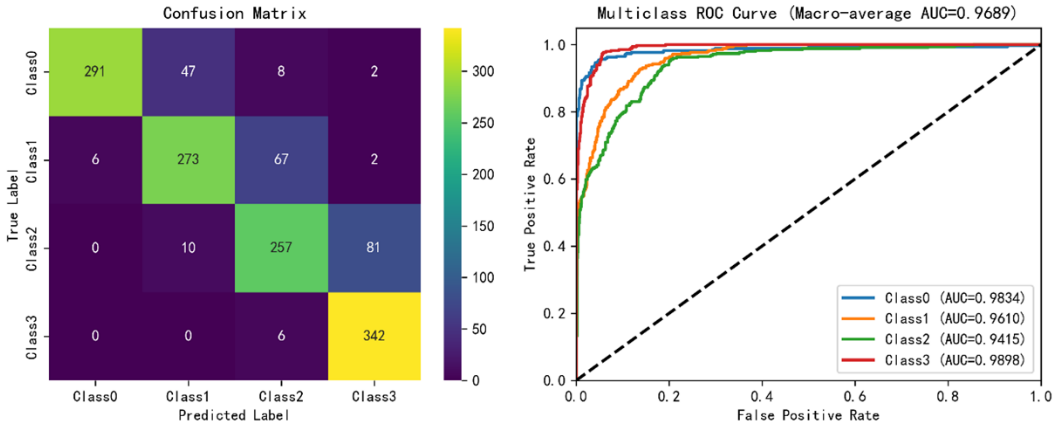


**Fig.S4. Performance evaluation of the baseline model on the challenge test set, including AUC-ROC curves and confusion matrix.**

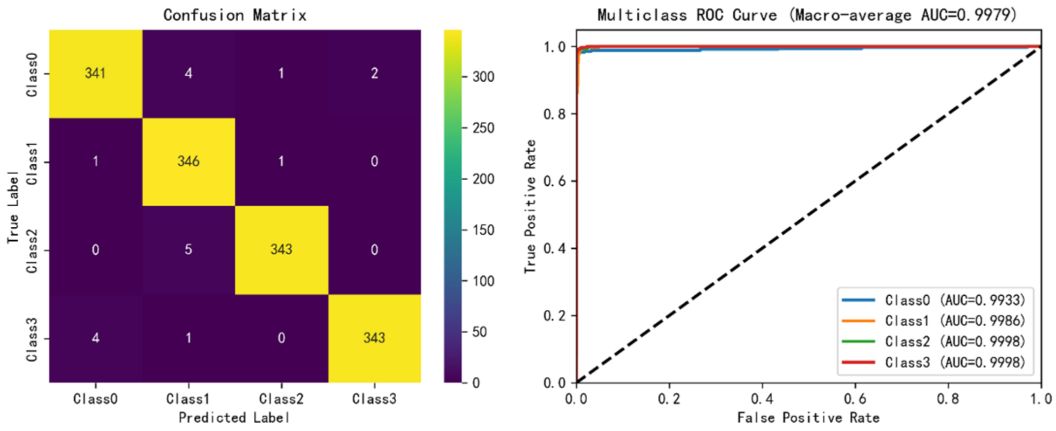


**Fig.S5. Performance evaluation of the mid model on the challenge test set, including AUC-ROC curves and confusion matrix.**

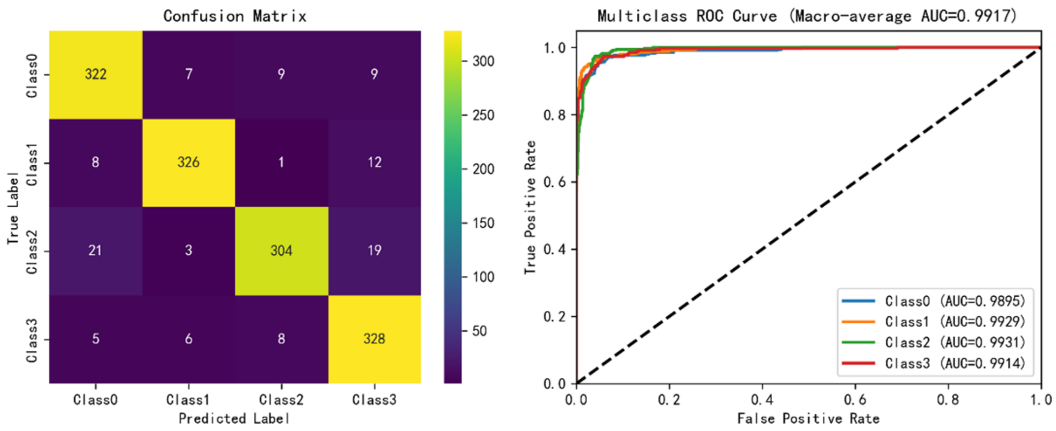


**Fig.S6. Performance evaluation of the full model on the challenge test set, including AUC-ROC curves and confusion matrix.**

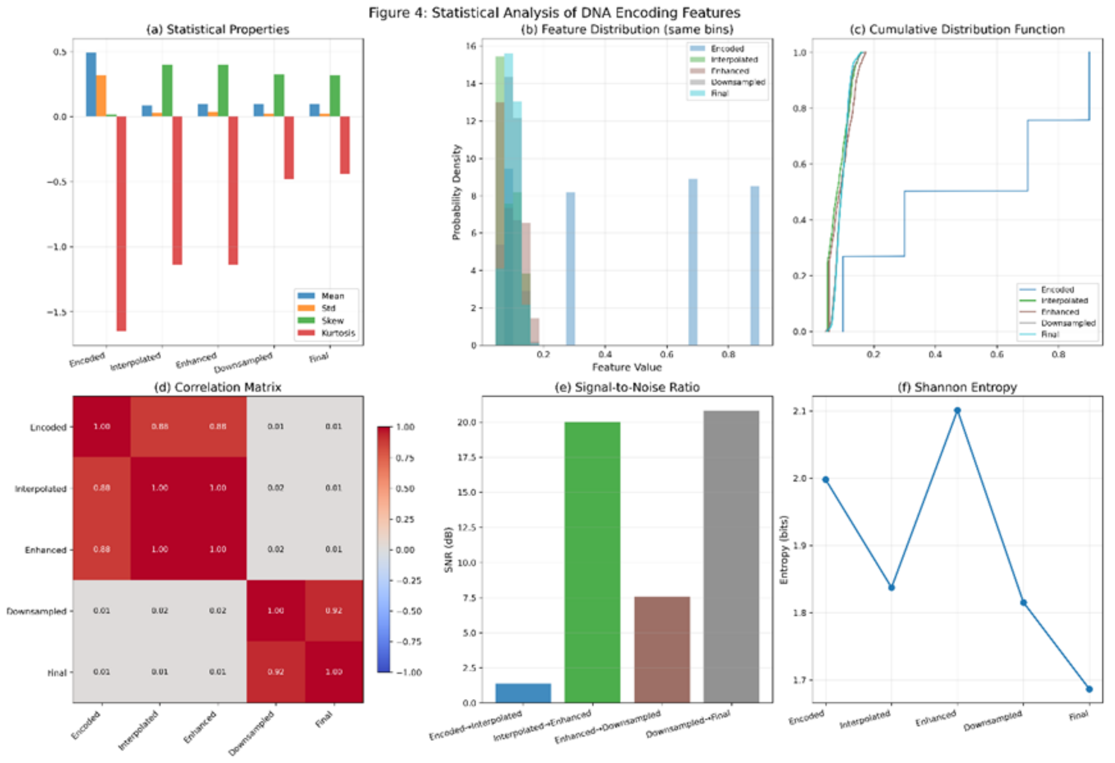


**Fig. S7. Visualization of learned features.** This figure illustrates the evolution of statistical properties of DNA-encoded signals throughout the complete UniWave pipeline, including encoding, interpolation, augmentation, compression, and filtering. (a) In terms of statistical metrics, the encoding stage exhibits low dispersion due to the discrete nature of nucleotide symbols. Interpolation and augmentation substantially increase the standard deviation and kurtosis, driving the features toward a continuous and highly diverse representation, whereas compression and filtering inversely reduce dispersion, causing the final signal to converge toward more compact feature distributions. (b) Feature distributions and (c) cumulative distribution functions (CDFs) further corroborate this trend: the encoded signals exhibit stepwise discrete distributions and CDFs, which become smooth and continuous after interpolation and augmentation, and subsequently reconverge toward central values following compression and filtering. (d) The correlation matrices indicate that the encoding, interpolation, and augmentation stages maintain correlation coefficients close to 1, preserving the core characteristics of the original signal, whereas the compression and filtering stages exhibit near-zero correlations with earlier stages, reflecting effective feature reconstruction. (e) Signal-to-noise ratio (SNR) varies across stages: interpolation introduces substantial modifications to the original signal, resulting in lower SNR, while augmentation and filtering act as fine-grained adjustments, yielding higher SNR. (f) Variations in Shannon entropy correspond to changes in information complexity, with the augmentation stage achieving the highest entropy (maximum information richness) and the final stage exhibiting the lowest entropy (most compact signal representation). Overall, this pipeline accomplishes an orderly transformation from discrete nucleotide signals to continuous features and ultimately to simplified representations. The stage-wise statistical variations are highly consistent with their respective operational objectives, preserving core information while effectively optimizing and simplifying the signal, thereby providing direct quantitative evidence for the scientific validity and effectiveness of the UniWave encoding framework.

In summary, this supplementary figure provides multi-dimensional quantitative validation of the core design advantages and practical engineering utility of the UniWave encoding framework. As a dedicated processing pipeline tailored for discrete DNA nucleotide signals, UniWave innovatively establishes a three-stage feature transformation paradigm—“discrete → continuous → simplified”—thereby overcoming the limitations of conventional DNA encoding methods, which are often constrained by low representational dimensionality and poor task adaptability. The central innovation lies in the combined use of interpolation and augmentation to achieve continuous lifting of discrete signals into higher-dimensional representations, while strong inter-stage correlations ensure faithful preservation of core nucleotide information. Subsequently, compression and filtering enable precise feature reconstruction, wherein weak correlations with earlier stages facilitate effective removal of redundant information. From a technical standpoint, parameter variations across stages do not constitute random signal perturbations; rather, they are tightly coupled to the predefined objectives of signal expansion, information enrichment, and feature purification. The stage-wise fluctuations in signal-to-noise ratio and Shannon entropy further substantiate the pipeline's precise regulatory capability in balancing information preservation and feature optimization. Compared with conventional encoding approaches, the distinctive value of UniWave lies in its ability to retain the intrinsic characteristics of DNA nucleotide signals while producing concise continuous features that are well suited for downstream tasks, thereby offering a standardized solution with clearly defined objectives and controllable performance for DNA information processing.

### Supplementary Tables：

Table S1. Data augmentation parameters across training and testing stages and their biological relevance analysis.

| Enhancement Stage | Enhancement Type | Technical Parameters | Biological Significance |
|---|---|---|---|
| Training Set Routine | Random Amplitude Scaling | Scaling Range: 0.8-1.2 (uniform)<br>Probability: 60% | Simulate signal intensity changes at different sequencing depths. |
| | Gaussian Noise Injection | Standard Deviation: 0.05 (SNR=26dB)<br>Probability: 80% | Simulate random errors in PCR amplification. |
| | Random Time Shift | Maximum Shift: ±60 bp<br>Probability: 40% | Simulate the uncertainty of read alignment and localization. |
| | Time Series Reversal | Reversal Probability: 30% | Simulate complementary strand sequencing scenarios. |
| Challenge Test | High-Frequency Noise | Standard Deviation: 0.10 (SNR=20dB) | Simulate sequencing noise in severely degraded samples. |
| | Random Time Reversal | Reversal Probability: 40% | Test the robustness of the model to sequence directionality. |
| | Random Local Occlusion | Occlusion Length: 100 - 300 bp<br>Position: 200 - 1600 bp<br>Probability: 60% | Simulate low-coverage regions in sequencing. |

Note: Some augmentation parameters are dynamically adjusted based on sequence length.

Table S2. Comparison of intrinsic noise robustness among different encoding strategies under a non-augmented training protocol.

| Encoding Stage | Model Params (K) | Feature Dimension | Standard Test Set | Challenge Test Set |
|---|---|---|---|---|
| Baseline Model | 25.6 | 1×1000 | BACC: 99.50% (95% CI: 0.9914-0.9985)<br>F1-macro: 0.9950<br>AUC-ROC: 0.9985 | BACC: 66.31% (95% CI: 0.6419-0.6845)<br>F1-macro: 0.6548<br>AUC-ROC: 0.8724 |
| Intermediate Model | 33.9 | 1×6000 | BACC: 99.28% (95% CI: 0.9882-0.9965)<br>F1-macro: 0.9928<br>AUC-ROC: 0.9980 | BACC: 66.16% (95% CI: 0.6426-0.6811)<br>F1-macro: 0.6418<br>AUC-ROC: 0.8934 |
| Full Model | 27.2 | 1×2000 | BACC: 96.76% (95% CI: 0.9578-0.9768)<br>F1-macro: 0.9677<br>AUC-ROC: 0.9957 | BACC: 72.98% (95% CI: 0.7048-0.7522)<br>F1-macro: 0.7283<br>AUC-ROC: 0.9200 |

Table S3. Evaluation of model robustness under a two-stage augmentation framework.

| Encoding Stage | Model Params (K) | Feature Dimension | Standard Test Set | Challenge Test Set |
|---|---|---|---|---|
| Baseline Model | 25.6 | 1×1000 | BACC: 98.20% (95% CI: 0.9752-0.9886)<br>F1-macro: 0.9820<br>AUC-ROC: 0.9965 | BACC: 83.55% (95% CI: 0.8172-0.8546)<br>F1-macro: 0.8349<br>AUC-ROC: 0.9689 |
| Intermediate Model | 33.9 | 1×6000 | BACC: 99.28% (95% CI: 0.9878-0.9965)<br>F1-macro: 0.9928<br>AUC-ROC: 0.9983 | BACC: 98.64% (95% CI: 0.9799-0.9923)<br>F1-macro: 0.9864<br>AUC-ROC: 0.9979 |
| Full Model | 27.2 | 1×2000 | BACC: 95.39% (95% CI: 0.9423-0.9645)<br>F1-macro: 0.9539<br>AUC-ROC: 0.9967 | BACC: 92.22% (95% CI: 0.9082-0.9358)<br>F1-macro: 0.9222<br>AUC-ROC: 0.9917 |

Table S4. Ablation study of key components within the complete waveform encoding architecture.

| Encoding Stage | Model Params (K) | Standard Test Set | Challenge Test Set | Reduction (Std, Chall) |
|---|---|---|---|---|
| Full Model | 27.2 | BACC: 95.39% (95% CI: 0.9423-0.9645)<br>F1-macro: 0.9539<br>AUC-ROC: 0.9967 | BACC: 92.22% (95% CI: 0.9082-0.9358)<br>F1-macro: 0.9222<br>AUC-ROC: 0.9917 | - |
| Remove WaveEncoder | 26.8 | BACC: 94.16% (95% CI: 0.9291-0.9534)<br>F1-macro: 0.9417<br>AUC-ROC: 0.9948 | BACC: 90.20% (95% CI: 0.8863-0.9178)<br>F1-macro: 0.9020<br>AUC-ROC: 0.9867 | 1.23%, 2.02% |
| Remove SelfAttention | 25.9 | BACC: 95.97% (95% CI: 0.9490-0.9695)<br>F1-macro: 0.9596<br>AUC-ROC: 0.9959 | BACC: 92.29% (95% CI: 0.9089-0.9361)<br>F1-macro: 0.9863<br>AUC-ROC: 0.9974 | -0.58%, -0.07% |
| Remove channel_attention | 26.1 | BACC: 95.24% (95% CI: 0.9411-0.9639)<br>F1-macro: 0.9524<br>AUC-ROC: 0.9963 | BACC: 91.43% (95% CI: 0.9003-0.9293)<br>F1-macro: 0.9147<br>AUC-ROC: 0.9878 | 0.15%, 0.79% |
| Remove Both SelfAttention & channel_attention | 24.8 | BACC: 96.61% (95% CI: 0.9562-0.9749)<br>F1-macro: 0.9661<br>AUC-ROC: 0.9969 | BACC: 86.60% (95% CI: 0.8475-0.8842)<br>F1-macro: 0.8668<br>AUC-ROC: 0.9721 | -1.22%, 5.62% |

Table S5. Performance comparison with conventional encoding methods under standard and challenge testing conditions.

| Encoding Stage | Model Params (K) | Feature Dimension | Standard Test Set | Challenge Test Set |
|---|---|---|---|---|
| Waveform Encoding | 27.2 | 1×2000 | BACC: 95.39% (95% CI: 0.9423-0.9645)<br>F1-macro: 0.9539<br>AUC-ROC: 0.9967 | BACC: 92.22% (95% CI: 0.9082-0.9358)<br>F1-macro: 0.9222<br>AUC-ROC: 0.9917 |
| One - hot Encoding | 26.9 | 4×1000 | BACC: 98.92% (95% CI: 0.9835-0.9943)<br>F1-macro: 0.9892<br>AUC-ROC: 0.9982 | BACC: 77.51% (95% CI: 0.7534-0.7981)<br>F1-macro: 0.7778<br>AUC-ROC: 0.9563 |
| Embedding Encoding | 1044.3 | 512×1000 | BACC: 99.35% (95% CI: 0.9886-0.9972)<br>F1-macro: 0.9935<br>AUC-ROC: 0.9976 | BACC: 72.41% (95% CI: 0.7036-0.7458)<br>F1-macro: 0.7416<br>AUC-ROC: 0.9805 |

Table S6. Cross-species generalization capability of waveform encoding in bacterial genome classification.

| Encoding Stage | Model Params (K) | Feature Dimension | Standard Test Set | Challenge Test Set |
|---|---|---|---|---|
| Full Model | 27.0 | 1×2000 | BACC: 93.61% (95% CI: 0.9158-0.9552)<br>F1-macro: 0.9361<br>AUC-ROC: 0.9818 | BACC: 92.34% (95% CI: 0.8988-0.9452)<br>F1-macro: 0.9233<br>AUC-ROC: 0.9758 |

Table S7. Experimental validation of numerical mapping.

| Nucleotide Order | Result | |
|---|---|---|
| | ACC | AUC-ROC |
| ATCG, GCTA | **0.6597 (95% CI: 0.6501~0.6700)** | **0.7195** |
| ATGC, CGTA | **0.6576 (95% CI: 0.6477~0.6669)** | **0.7107** |
| ACTG, GTCA | 0.6144 (95% CI: 0.6044~0.6247) | 0.6506 |
| ACGT, TGCA | 0.6230 (95% CI: 0.6131~0.6339) | 0.6660 |
| AGTC, CTGA | 0.6328 (95% CI: 0.6226~0.6425) | 0.6813 |
| AGCT, TCGA | 0.6404 (95% CI: 0.6310~0.6510) | 0.6838 |
| TACG, GCAT | **0.6626 (95% CI: 0.6530~0.6721)** | **0.7164** |
| TAGC, CGAT | **0.6522 (95% CI: 0.6423~0.6617)** | **0.7050** |
| TCAG, GACT | 0.6166 (95% CI: 0.6062~0.6264) | 0.6662 |
| TGAC, CAGT | 0.6042 (95% CI: 0.5939~0.6144) | 0.6601 |
| CATG, GTAC | 0.6362 (95% CI: 0.6261~0.6464) | 0.6845 |
| CTAG, GATC | 0.6431 (95% CI: 0.6334~0.6520) | 0.7032 |

Note: GC and AT nucleotides inherently possess distinct chemical structures, and GC content is closely associated with functional regions and fragmentation patterns in many genomic analysis tasks. The above experiments indicate that the model achieves higher performance when separating the AT group from the GC group, suggesting that the model partially leverages the "GC preference" sequence statistical feature for classification. When the numerical encoding captures the intrinsic physicochemical properties of nucleotides, the model can more effectively learn the mapping between sequences and their functional attributes.
